# Effect of alginate encapsulation on growth and viability of polycyclic aromatic hydrocarbon-degrading bacteria varies by environment, species, and capsule design

**DOI:** 10.64898/2026.08.26.747349

**Authors:** Amelia M. Foley, Claudia K. Gunsch

## Abstract

Polycyclic aromatic hydrocarbons (PAHs) are hazardous organic contaminants for which microbial bioaugmentation is a promising remediation strategy, but poor persistence of introduced microorganisms can limit efficacy. Encapsulation may improve persistence, yet the influence of capsule design, microbial species, and environmental conditions on performance remains poorly understood. We evaluated alginate encapsulation of the PAH-degrading bacteria *Pseudomonas putida* and *Novosphingobium aromaticivorans* across nutrient conditions and capsule formulations. Encapsulation effects varied by species and medium, influencing growth rate, maximum cell density, overall growth, and lag time; notably, encapsulation shortened lag time of *N. aromaticivorans* in sRB_15_ medium (36.9 h to 3.9-5.3 h). Enumeration methods also affected apparent cell recovery. After 8 weeks, encapsulation had no significant effect on *P. putida* but resulted in increased concentrations of *N. aromaticivorans* relative to planktonic cultures (1.22 x 10^8^ CFU/mL vs. 2.05 x 10^6^ CFU/mL). Capsule composition further influenced cell retention: increasing alginate approximately doubled capsule-associated cell concentrations, while chitosan coatings reduced cell concentrations within capsules without affecting external concentrations. These findings demonstrate that the benefits of encapsulation are species- and environment-dependent and that capsule formulation can be tuned to influence bacterial persistence and release, informing the design of encapsulated inoculants for bioaugmentation applications.

## Introduction

Bioaugmentation involves the controlled addition of microorganisms with specific functional capabilities to an environment to promote desired outcomes such as contaminant degradation, agricultural productivity, and wastewater treatment. However, successful bioaugmentation requires introduced microorganisms to remain viable and functionally active following inoculation. Exogenous strains often struggle to establish in new environments due to competition, environmental conditions, and other factors contributing to colonization resistance (M. Wu et al., 2016; Yu et al., 2005). Consequently, the transience of introduced strains is often cited as a cause of the failure of these approaches, especially for in-situ bioremediation in harsh conditions such as polycyclic aromatic hydrocarbon (PAH) contaminated environments (Haleyur et al., 2019; Yu et al., 2005). These limitations suggest that additional strategies are required to promote the persistence of introduced microorganisms.

Encapsulation offers one strategy to improve microbial delivery and persistence by immobilizing cells within a protective polymer matrix (Bayat et al., 2015; Partovinia & Rasekh, 2018). Encapsulation can protect cells from environmental stressors such as shear damage, pH, temperature, predation, and hazardous compounds. Capsules provide space for microbial growth and controlled release of cells (Cassidy et al., 1996; Perullini et al., 2015). Among available encapsulation methods, ionic gelation with sodium alginate is widely used because it occurs under mild conditions suitable for cell survival. Sodium alginate is easy to use, inexpensive, nontoxic, and biodegradable (Valdivia-Rivera et al., 2021). Chitosan, another natural polymer, is commonly used as a coating for alginate capsules to increase capsule stability by forming a polycation membrane (Gaserod et al., 1998).

While encapsulation is often treated as a generally beneficial delivery strategy, its effectiveness is context dependent. Changes in capsule properties such as polymer concentration, coating, and cell density can greatly influence bacterial growth within capsules, release of cells, mass transfer, and contaminant degradation (Essifi et al., 2021; Faria et al., 2021; He et al., 2015; Z. Wu et al., 2012). However, studies evaluating the relationship between specific aspects of capsule design and microbial outcomes are still limited (Lobel et al., 2024).

Importantly, encapsulation performance also depends on characteristics of both the microorganism and its environment. Despite the potential of alginate encapsulation, its ability to enhance bacterial growth and bioremediation performance has been variable, depending on strain selection and environmental conditions (Errampalli et al., 1999; Paje et al., 1998). Environmental parameters such as nutrient availability can influence the survival of encapsulated bacteria and their degradation of pollutants. Similarly, microorganisms with different physiological and ecological characteristics may respond differently to encapsulation. Previous studies have often focused on encapsulating individual strains in isolation (Dou et al., 2021; Huizenga et al., 2024; Q. Liu et al., 2023; Lu et al., 2020; Park et al., 2021; Ren et al., 2022; Song et al., 2021; Wang et al., 2023), leaving the effect of encapsulation on different types of microbes underexplored. Understanding how capsule design interacts with microbial characteristics and environmental conditions is therefore important for rationally designing encapsulated inoculants for different bioremediation applications.

Quantifying these interactions presents an additional challenge because cells entrapped within a polymer matrix cannot be readily enumerated. To quantify live immobilized biomass, cells typically must be released via decapsulation. Sodium citrate is commonly used to reverse alginate crosslinking, but decapsulation procedures may affect cell recovery and consequently bias estimates of viability (Champagne et al., 2011). Alternative approaches, including optical density and fluorescence-based measurements, may complement culture-based enumeration but require validation for encapsulated systems.

In this study, we hypothesized that the effects of alginate encapsulation on bacterial growth, persistence, and release would depend on capsule design, bacterial species, and nutrient environment. We compared the growth and long-term viability of encapsulated *Pseudomonas putida* and *Novosphingobium aromaticivorans* with their planktonic counterparts to assess the ability of encapsulation to protect PAH-degrading bacteria for bioremediation applications. We investigated alginate capsule designs, including 1% and 2% alginate, a chitosan coating, and different initial cell densities. We also applied complementary approaches for estimating microbial concentrations, including optical density, decapsulation followed by plate counts, and fluorescence intensity, to distinguish short-term growth effects from long-term persistence and cell retention/release.

## Methods

### Chemicals

For bacterial culture preparation, Luria-Bertani (LB) broth, LB agar, and R2A broth were obtained from Fisher Scientific. For encapsulation experiments, sodium alginate was obtained from VWR. Calcium chloride dihydrate, sodium citrate, and chitosan (low molecular weight) were obtained from Sigma Aldrich.

### Strains and media

The strains *Pseudomonas putida* PVP102 (mScarlet-I, mVenus, kan^R^, cam^R^) and *Novosphingobium aromaticivorans* PMN120 (mScarlet-I, GFPmut2, kan^R^) were selected in this work for their PAH-degrading ability, constitutive fluorescence, and difference in 16S rRNA copy number (Varner et al., 2022). *P. putida* and *N. aromaticivorans* were grown from glycerol stocks at 30°C on LB and R2A medium, respectively. Liquid sRB_15_ medium for experimental reactors was prepared as previously described, with 0.2% pyruvate (Volkoff et al., 2022).

### Encapsulation

Cells were encapsulated in sodium alginate microcapsules using the ionic gelation technique (Yeung et al., 2016). Briefly, 1.5 x 10^8^ bacterial cells were concentrated by centrifugation, then resuspended in 150 μL LB medium and added to a 2% sodium alginate solution at a ratio of 1% v/v in 15 mL alginate for a final concentration of 1 x 10^7^ CFU/mL alginate. The mixture was gently inverted to mix and then transferred to sterile 20mL syringes using 16-gauge needles. Syringes were rested upright with headspace for 1 h to allow air bubbles to degas. The solution was extruded via 25-gauge needles into 250 mL of continuously stirred calcium chloride (0.1M) crosslinking solution with a syringe pump. Capsules were crosslinked in calcium chloride for 1 h before being rinsed three times with PBS (pH 7.4). Abiotic capsules were prepared following the same procedure without the addition of bacterial cells.

### Chitosan coating

0.5% w/v chitosan was prepared according to previous methods (Pupa et al., 2021; Yeung et al., 2016; Zhou et al., 1998). Briefly, 0.5 g chitosan was dissolved in 90 mL distilled water and 0.8 mL acetic acid. pH was adjusted to 5.0-5.1 with 1M NaOH, and the total volume was adjusted to ∼100 mL. The solution was autoclaved and filtered prior to use. To prepare coated capsules, rinsed alginate capsules were submerged in 100 mL 0.5 % w/v chitosan and mixed by shaking for 1 h at 100 rpm at 21°C. Capsules were then filtered and rinsed three times with ∼50 mL of sterile PBS (pH 7.4).

### Decapsulation and viability in sodium citrate

For planktonic groups, an overnight culture of *P. putida* was prepared and adjusted to an OD_600_ of 1. Then, 200 μL of the *P. putida* culture was aliquoted to microcentrifuge tubes. 1000 μL of 0.1 M sodium citrate or 1000 μL 1x PBS was added to tubes in triplicate. Samples were immediately taken at the 0-, 15-, and 30-minute mark. Samples were serially diluted and plated on LB agar to quantify bacterial concentrations in samples. For encapsulated groups, 2% alginate *P. putida* capsules were prepared as previously described. 200 μL capsules were placed in microcentrifuge tubes and 1000 μL sodium citrate was added to tubes in triplicate. Supernatant samples were taken at the 0-, 15-, and 30-minute time points. Tubes were vortexed occasionally throughout the wait step. Samples were serially diluted and plated to quantify bacterial concentrations.

### Viability experiment

Experiments testing cellular viability were conducted in disposable borosilicate glass culture tubes (16×100mm, VWR). 2 g of capsules (wet weight) containing ∼2 x 10^7^ cells (calculated based on the density of alginate capsules being ∼1 g/mL) were placed in each reactor in 10 mL of sRB_15_ media containing pyruvate. In order to ensure equal numbers of cells in both encapsulated and planktonic conditions, ∼2 x 10^7^ cells in 20 μL LB were added to planktonic reactors. Reactors were incubated shaking (100 rpm) at room temperature for 8 weeks. Reactors were prepared in triplicate and were sacrificed at each time point. Experiments were performed in batches to evaluate one capsule formulation variable at a time (Table 1). Capsules were prepared using *P. putida*, 2% alginate, no coating, and 10^7^ colony-forming units per milliliter (CFU/mL) cell loading density unless otherwise noted.

**Table 1.**
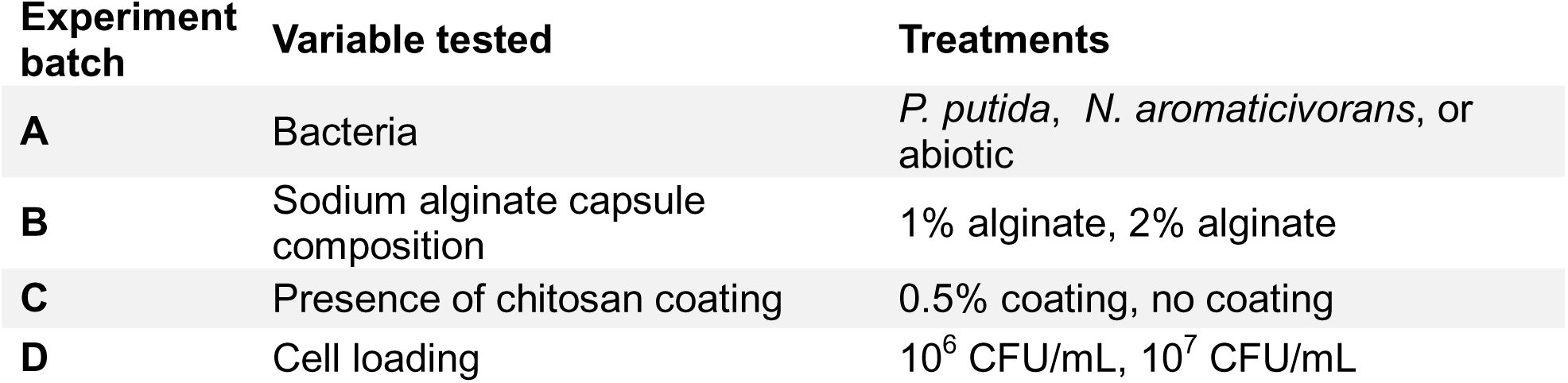
Experimental design for 8-week viability assessment.

Reactors were sampled on days 0, 14, 28, 42, and 56. Capsules were collected using a sterile spatula and their volume was measured in a demarcated 1.7 mL microcentrifuge tube. To dissolve capsules, 1000 μL sodium citrate was added to 200 μL capsules and incubated at room temperature for 15 minutes with occasional vortexing. Supernatant (200 μL), microcapsules (200 μL), and dissolved microcapsules (200 μL) were collected in duplicate and placed in a 96 well black polystyrene microplate (flat bottom, clear)(Corning). OD_600_ and fluorescence measurements were taken on a Tecan Infinite M1000 Pro plate reader. Fluorescence intensity was measured in top mode for mScarlet-I (ex: 569nm, em: 593nm), mVenus (ex: 500nm, em: 545nm), and GFPmut2 (ex: 475nm, em: 515nm) at a constant manual gain for each fluorophore across timepoints. Multiple reads per well were averaged (filled square, 4×4) and z-position was calculated from wells. A random subset of samples was enumerated by plate count at each intermediate time point. On the final day, all samples were enumerated by plate count using the drop plate method (Thomas et al., 2015). For plate counts, capsules were dissolved by adding 1000 μL sodium citrate (0.1 M) to 200 μL of microcapsules for 15 minutes (vortexing occasionally).

### Growth curves

Bacteria, media, and capsules were prepared as previously described. 0.2 g capsules were added to each well of a 48-well plate, in 1000 μL media. For planktonic wells, an equivalent number of bacterial cells were inoculated (∼2 x 10^6^ cells in 2 μL LB). The well plate was incubated shaking (orbital, 3mm amplitude) at 30°C for 60 – 90 h and absorbance measurements at 600 nm were taken every 30 minutes with multiple reads per well (filled circle, 7×7). Capsules were evaluated in both LB and sRB_15_ media. Abiotic capsules were included on each plate as a blank control. Measured OD_600_ values were corrected based on blank measurements.

### Imaging

The appearance of *P. putida* within alginate capsules was characterized with a confocal microscope. Capsules were prepared as previously described, incubated for 24 hours in LB at 30°C, then washed in PBS and stored overnight in PBS at 4°C. Capsules were placed in a glass-bottom 48 well plate for imaging. Images were acquired with a Zeiss LSM 900 confocal microscope (Zeiss | Oberkochen, Germany). Images were recorded as z-stacks with the 10x and 40x objectives and processed in Fiji for coloring and contrast.

### Data analysis

Data were analyzed using RStudio, version 2024.09.0. Experimental replicates for all data were reported as averages and standard error. The package gcplyr was used to analyze growth curve data and calculate growth rate (h^-1^), maximum cell density (OD_600_), area under the curve (AUC)(OD_600_*h), and lag time (h) (Blazanin, 2024). Growth phases were identified in gcplyr by locating local minima/maxima in smoothed derivative curves (see Supplemental Material for greater detail). Total encapsulated reactor concentration was calculated using the concentrations and volumes of each component (capsule and supernatant).

We performed multiple linear regression analyses to assess the relationship between *P. putida* bacterial concentration and the variables time, treatment, and fluorescence. The analysis was conducted using lm() in the stats package in R. Prior to analysis, a log10 transformation was applied to bacterial concentration to normalize the data. The Akaike Information Criterion (AIC) was calculated for models using the AIC() function to determine the relative quality of the models by measuring goodness of fit and complexity (Akaike, 1974). All other results were analyzed using ANOVA followed by post hoc Tukey’s HSD test. Statistical significance was determined with a 95% confidence interval.

## Results

### Growth of tested strains under encapsulated and planktonic conditions

The growth profiles of encapsulated *P. putida* and *N. aromaticivorans* and their planktonic counterparts were monitored over time in rich (LB) and minimal (sRB15) media by measuring OD_600_ (Fig. 1). Quantitative growth metrics are summarized in Table 2. For encapsulated groups, two distinct growth phases were observed: an initial phase of rapid growth (phase 1) followed by a secondary increase in OD_600_ (phase 2). Across both species, the effects of encapsulation on growth rate, maximum density, overall growth (area under the curve [AUC]), and lag time depended on the growth medium and bacterial species.

**Figure 1.**
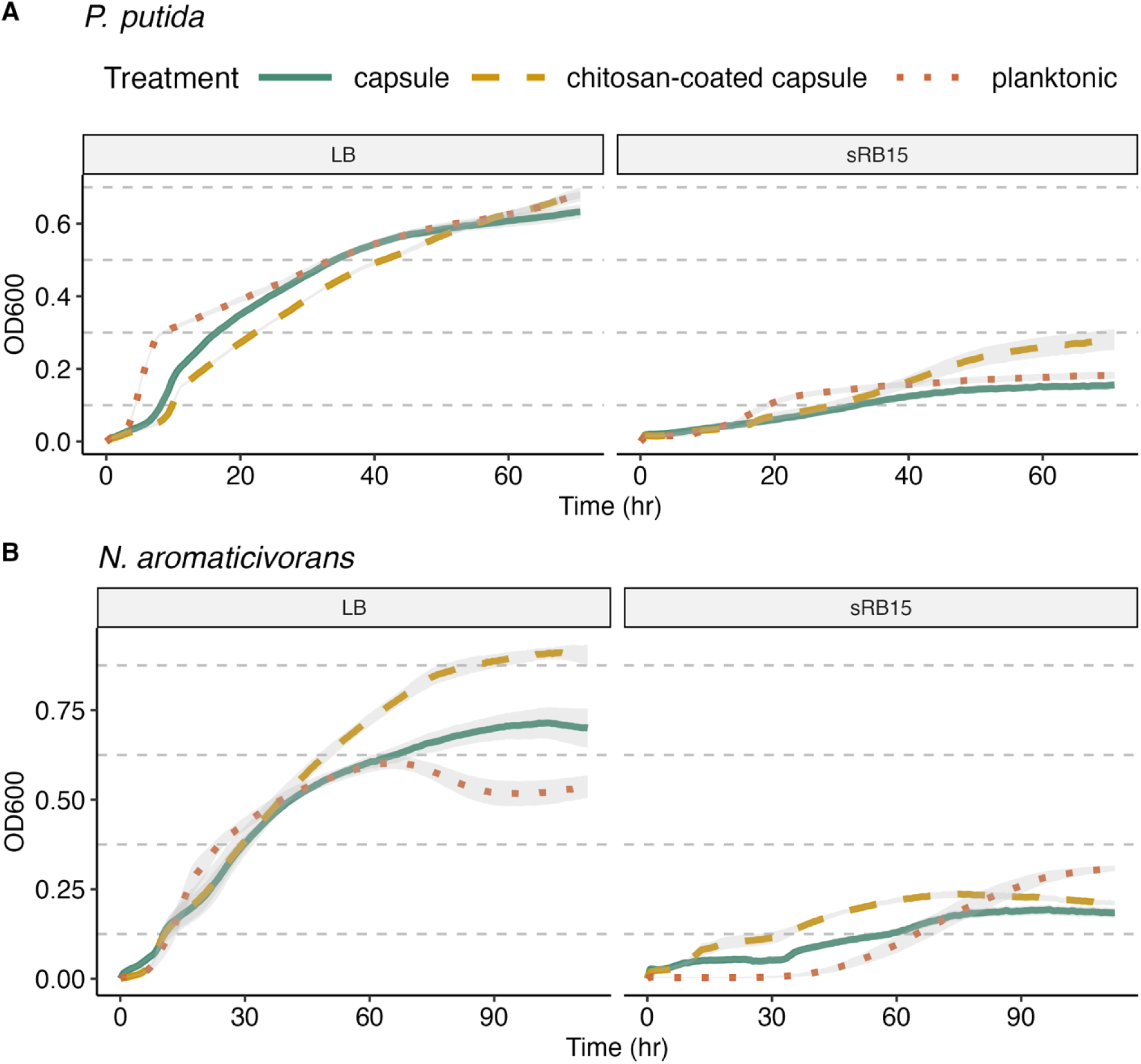
Growth curves for 3 treatment types in LB and sRB_15_ media for A) *P. putida* and B) *N. aromaticivorans*. Grey regions represent standard error (n > 8).

**Table 2.** Summary statistics for growth curve metrics calculated from data shown in Fig. 1. SE represents standard error. Means not sharing any letter are significantly different by the Tukey HSD test (p < .05). Comparisons are made only for treatments within the same strain/media groups.

| Strain | Media | Treatment | n | Growth<br>Rate ($\text{h}^{-1}$)<br>(Phase 1) | | Growth<br>Rate ($\text{h}^{-1}$)<br>(Phase 2) | | Density<br>($\text{OD}_{600}$) | | AUC<br>($\text{OD}_{600} \cdot \text{h}$) | | Lag Time (h)<br>(Phase 1) | |
| --- | --- | --- | --- | --- | --- | --- | --- | --- | --- | --- | --- | --- | --- |
|  |  |  |  | Mean | SE | Mean | SE | Mean | SE | Mean | SE | Mean | SE |
| *N. aroma* | LB | Capsule | 10 | 0.36<sup>a</sup> | 0.08 | 0.12<sup>a</sup> | 0.01 | 0.73<sup>a</sup> | 0.04 | 55.87<sup>a</sup> | 1.39 | 3.38<sup>a</sup> | 0.33 |
|  |  | Chitosan | 8 | 0.52<sup>a</sup> | 0.04 | 0.13<sup>a</sup> | 0.00 | 0.92<sup>b</sup> | 0.02 | 66.54<sup>b</sup> | 2.71 | 3.33<sup>a</sup> | 0.28 |
|  |  | Planktonic | 10 | 0.49<sup>a</sup> | 0.07 | — | — | 0.61<sup>c</sup> | 0.02 | 50.59<sup>a</sup> | 2.65 | 4.92<sup>a</sup> | 0.89 |
| *N. aroma* | sRB<sub>15</sub> | Capsule | 11 | 0.12<sup>a</sup> | 0.01 | 0.14<sup>a</sup> | 0.05 | 0.20<sup>a</sup> | 0.01 | 13.51<sup>a</sup> | 0.96 | 3.89<sup>a</sup> | 0.30 |
|  |  | Chitosan | 8 | 0.16<sup>a</sup> | 0.01 | 0.09<sup>a</sup> | 0.00 | 0.24<sup>a</sup> | 0.01 | 19.14<sup>b</sup> | 1.10 | 5.29<sup>a</sup> | 0.07 |
|  |  | Planktonic | 11 | 0.34<sup>b</sup> | 0.05 | — | — | 0.31<sup>b</sup> | 0.01 | 13.17<sup>a</sup> | 1.38 | 36.94<sup>b</sup> | 3.77 |
| *P. putida* | LB | Capsule | 10 | 0.32<sup>a</sup> | 0.04 | 0.36<sup>a</sup> | 0.02 | 0.65<sup>a</sup> | 0.02 | 29.98<sup>a</sup> | 0.57 | 2.58<sup>a</sup> | 0.01 |
|  |  | Chitosan | 8 | 0.33<sup>a</sup> | 0.05 | 0.33<sup>a</sup> | 0.04 | 0.68<sup>a</sup> | 0.01 | 27.56<sup>b</sup> | 0.36 | 2.57<sup>a</sup> | 0.01 |
|  |  | Planktonic | 10 | 0.64<sup>b</sup> | 0.06 | — | — | 0.70<sup>a</sup> | 0.02 | 32.48<sup>c</sup> | 0.49 | 2.55<sup>a</sup> | 0.01 |
| *P. putida* | sRB<sub>15</sub> | Capsule | 11 | 0.23<sup>a</sup> | 0.06 | 0.04<sup>a</sup> | 0.00 | 0.16<sup>a</sup> | 0.01 | 6.87<sup>a</sup> | 0.61 | 5.06<sup>a</sup> | 1.32 |
|  |  | Chitosan | 8 | 0.18<sup>a</sup> | 0.05 | 0.12<sup>b</sup> | 0.02 | 0.28<sup>b</sup> | 0.03 | 9.82<sup>b</sup> | 1.03 | 3.09<sup>a</sup> | 0.10 |
|  |  | Planktonic | 11 | 0.27<sup>a</sup> | 0.07 | — | — | 0.19<sup>a</sup> | 0.01 | 8.78<sup>ab</sup> | 0.53 | 5.47<sup>a</sup> | 0.39 |

For *P. putida* in LB, planktonic cultures displayed a significantly higher phase 1 growth rate (0.64 h^-1^) compared with both encapsulated treatments (capsule = 0.32 h^-1^ chitosan = 0.33 h^-1^). Planktonic cultures also had greater overall growth than either encapsulated treatment, whereas maximum density and lag time did not differ among treatments. In sRB_15_, phase 1 growth rates were similar among treatments; however, chitosan-coated capsules produced a higher phase 2 growth rate than alginate capsules. Chitosan-coated capsules also resulted in a higher maximum density than both uncoated capsules and planktonic cultures. Overall growth was also greater for chitosan-coated than uncoated capsules, although neither encapsulated treatment differed significantly from the planktonic treatment. Lag time did not differ significantly among treatments.

In contrast, *N. aromaticivorans* showed stronger treatment-dependent responses to encapsulation. In LB, encapsulation increased maximum density, and chitosan-coated capsules significantly enhanced overall growth compared with both other treatments. Growth rates and lag times were unaffected by treatment in LB. In sRB_15_, encapsulation reduced the phase 1 growth rate and maximum density compared with the planktonic group. However, chitosan-coated encapsulation increased overall growth (AUC) compared with planktonic and alginate-encapsulated groups. Notably, encapsulation substantially shortened lag time, from 36.94 ± 3.77 h in planktonic cultures to 3.89 ± 0.30 and 5.29 ± 0.07 h in alginate and chitosan-coated capsules, respectively.

### Characterization and recovery of encapsulated *Pseudomonas putida*

When encapsulated in 2% alginate beads and incubated in LB for 24 h at 30°C, *P. putida* cells formed aggregates 10–20 μm in diameter distributed throughout the alginate capsule (Fig. 2A). mScarlet fluorescence revealed dense bacterial clusters within the capsule matrix, including clusters near the capsule surface (Fig. 2B).

**Figure 2.**
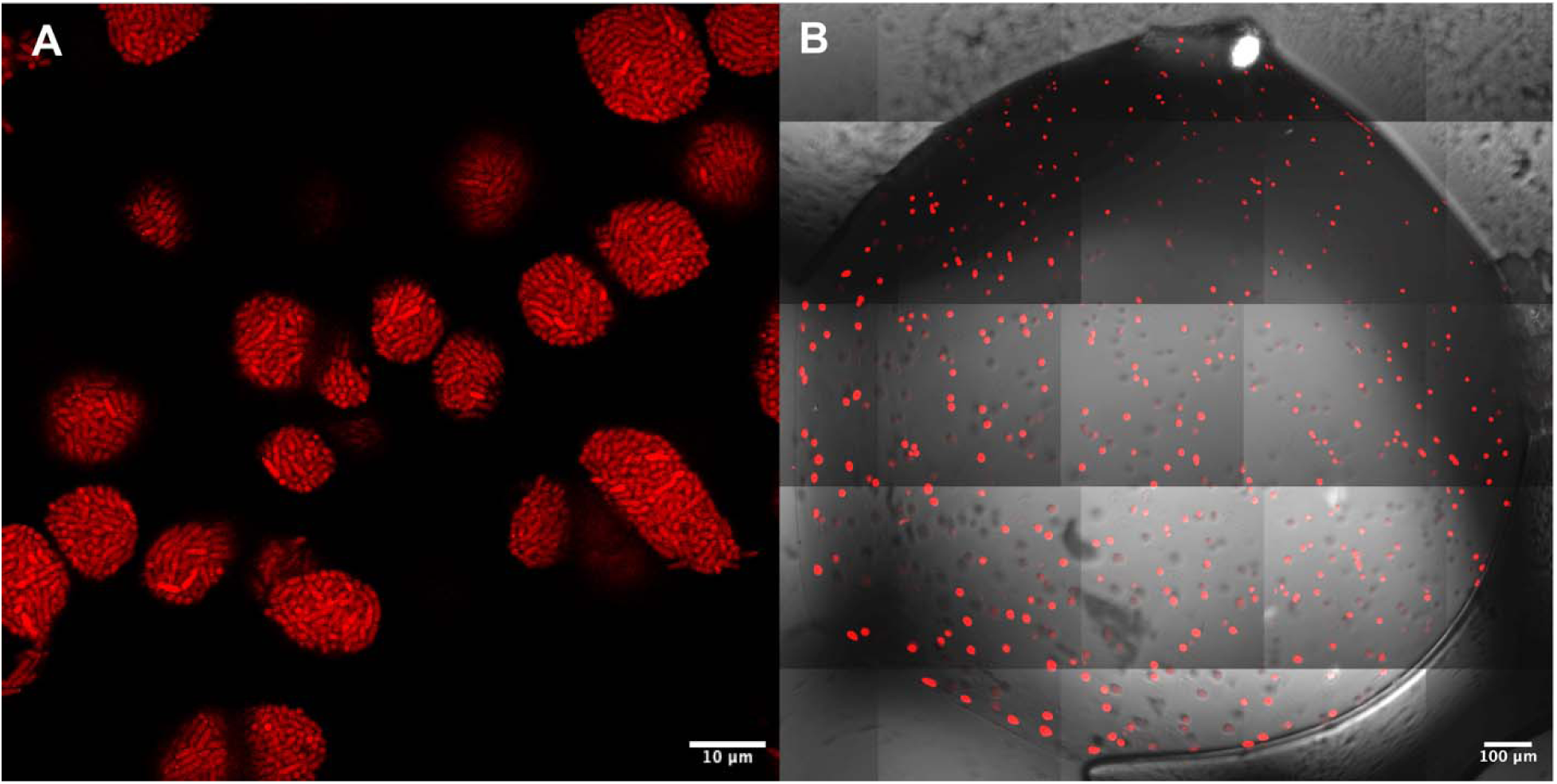
Confocal laser scanning microscopy images of encapsulated *P. putida*. A) Multiple pores containing bacterial cells within a singular alginate capsule. B) Confocal image overlayed with brightfield image of a singular *P. putida* microcapsule.

The effect of sodium citrate exposure on cell viability was evaluated for planktonic *P. putida* using PBS as a control. A 30-minute exposure to sodium citrate decreased recoverable viable cell concentrations compared with PBS (Fig. 3). For encapsulated *P. putida*, 5–15 minutes of sodium citrate exposure increased measured CFU/mL as capsules dissolved and released entrapped cells. After 30 minutes, however, recoverable CFU/mL decreased, consistent with an adverse effect of prolonged sodium citrate exposure on cell viability. Based on these results, a 15-minute decapsulation protocol was used in subsequent experiments to minimize the effect of sodium citrate on viability.

**Figure 3.**
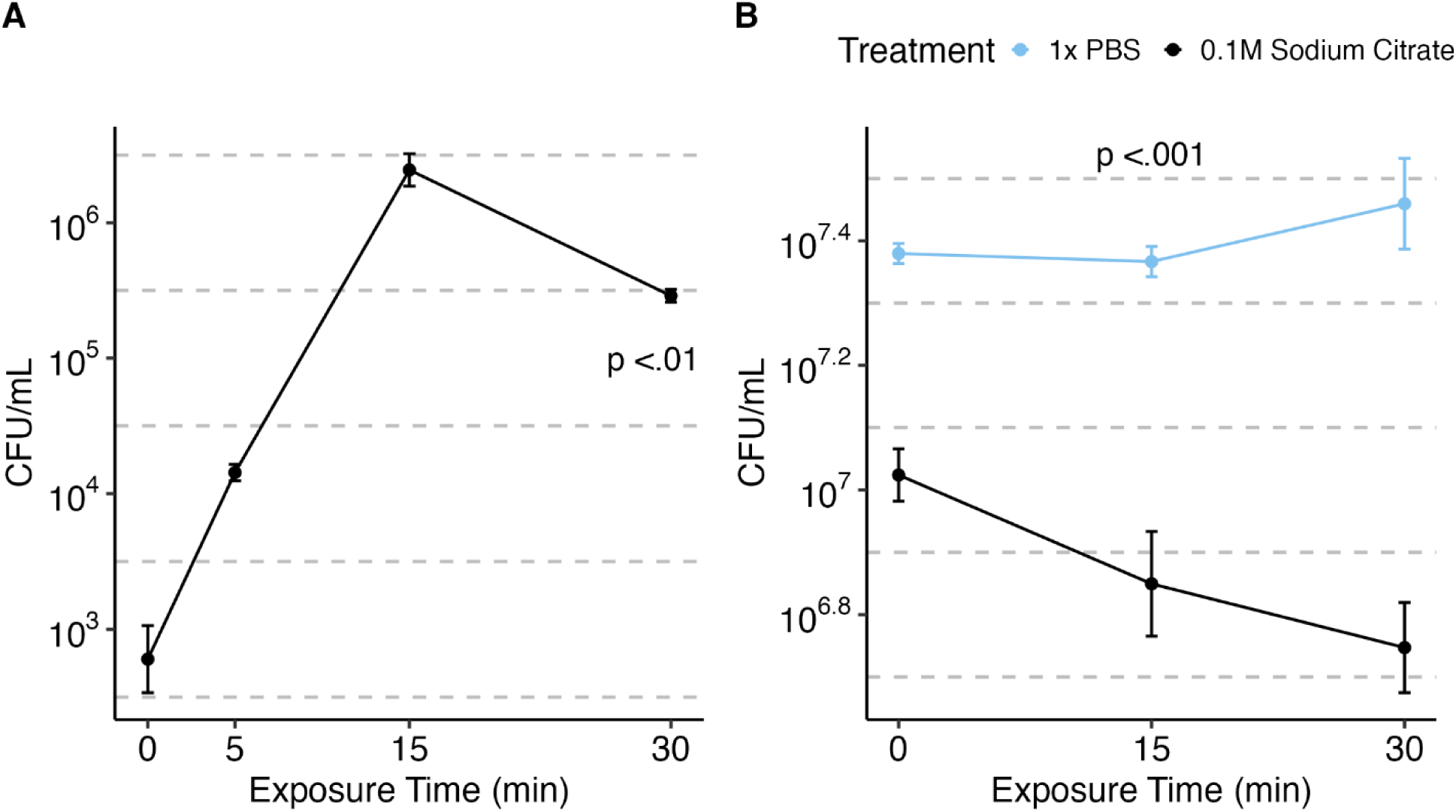
Viability of *P. putida* in sodium citrate for A) encapsulated cells and B) planktonic cells. Error bars represent standard error (n = 3). P-values refer to the effect of time (A) and the effect of treatment (B).

### Long term viability

After 8 weeks of incubation in sRB15 containing pyruvate, capsules maintained high cell concentrations of both *P. putida* and *N. aromaticivorans* (Fig. 4). Total cell concentrations in encapsulated reactors were calculated from the concentrations and volumes of the capsule and supernatant fractions to enable comparison with planktonic reactors. For *P. putida*, there was no significant difference between encapsulated and planktonic reactor concentrations. In contrast, total *N. aromaticivorans* concentrations were significantly higher in encapsulated than planktonic reactors (1.22 x 10^8^ CFU/mL vs. 2.05 x 10^6^ CFU/mL).

**Figure 4.**
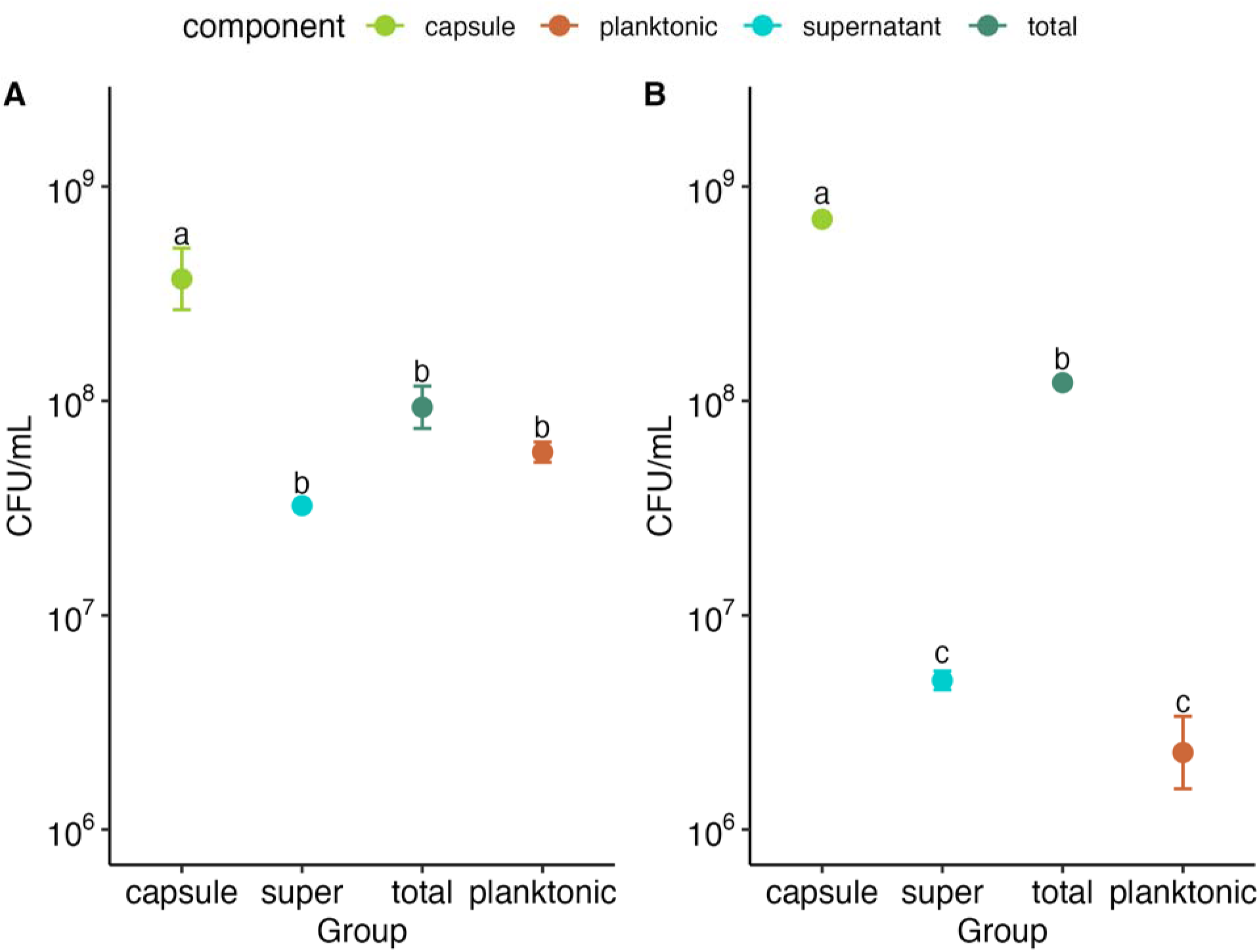
Viability/survival after 8-week incubation for A) *P. putida* and B) *N. aromaticivorans*. Error bars represent standard error (n = 3). Means not sharing any letter are significantly different by the Tukey HSD test (p < .05)

The effect of alginate concentration on long-term *P. putida* viability was evaluated by comparing 1% and 2% alginate capsules after 8 weeks in sRB15 containing pyruvate (Fig. 5). The 2% alginate capsules contained a higher mean cell concentration than 1% alginate (2.12 x 10^8^ versus 1.01 x 10^8^ CFU/mL. There was no significant difference in supernatant or total reactor concentrations between 1% and 2% alginate capsules, while planktonic concentrations remained higher than encapsulated reactor totals for both alginate formulations.

**Figure 5.**
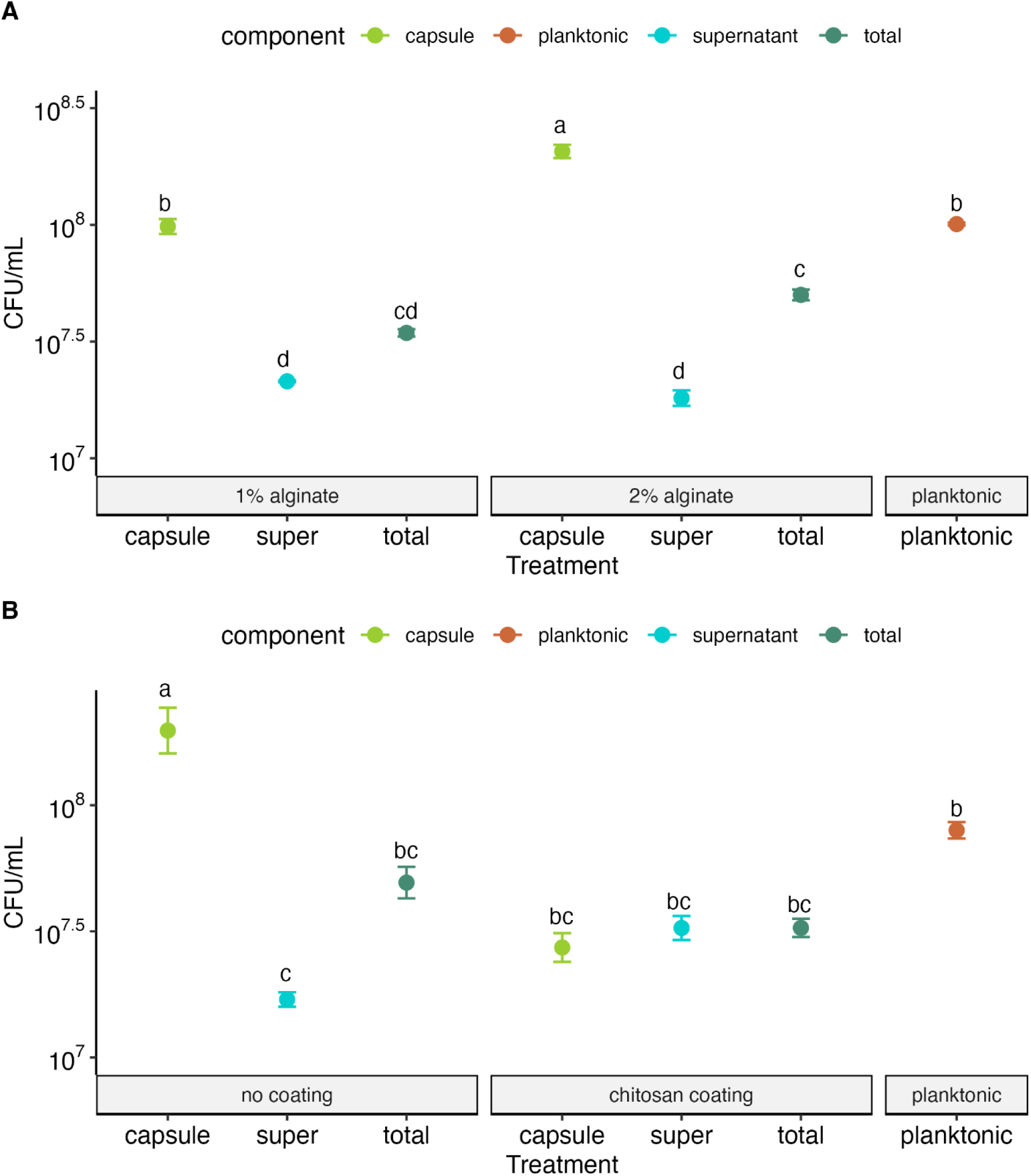
Viability/survival after 8-week incubation for *P. putida* capsules formulated with varying A) alginate concentration and B) chitosan coating status. Error bars represent standard error (n = 3). Means not sharing any letter are significantly different by the Tukey HSD test (p < .05)

The addition of a chitosan coating reduced the cell concentration within the capsule from 2.4 × 10^8^ to 2.98 × 10^7^ CFU/mL after 8 weeks (Fig. 5). However, no significant difference was observed in supernatant or total reactor concentrations between capsules with or without a chitosan coating. Total reactor concentrations also did not differ between planktonic and encapsulated treatments. Initial cell density (10^6^ vs. 10^7^ CFU/mL) did not significantly affect cell concentrations after 8 weeks (Fig. S1).

### Fluorescence as a proxy for concentration

We evaluated whether fluorescence could serve as a proxy for viable cell concentration in the fluorescently tagged strains. Standard curves demonstrated a strong linear relationship between CFU/mL and relative fluorescence units (RFU) (R^2^ = 0.97) for *P. putida*, which expressed mScarlet-I and mVenus (Fig. S2). We then evaluated this relationship using data from the 8-week experiment. The fluorescence of intact and dissolved capsules did not explain variability in capsule CFU/mL. For supernatant samples, the relationship between fluorescence and CFU/mL changed over time, with the relationship between RFU and cell concentration shifting across sampling days (Fig. 6).

**Figure 6.**
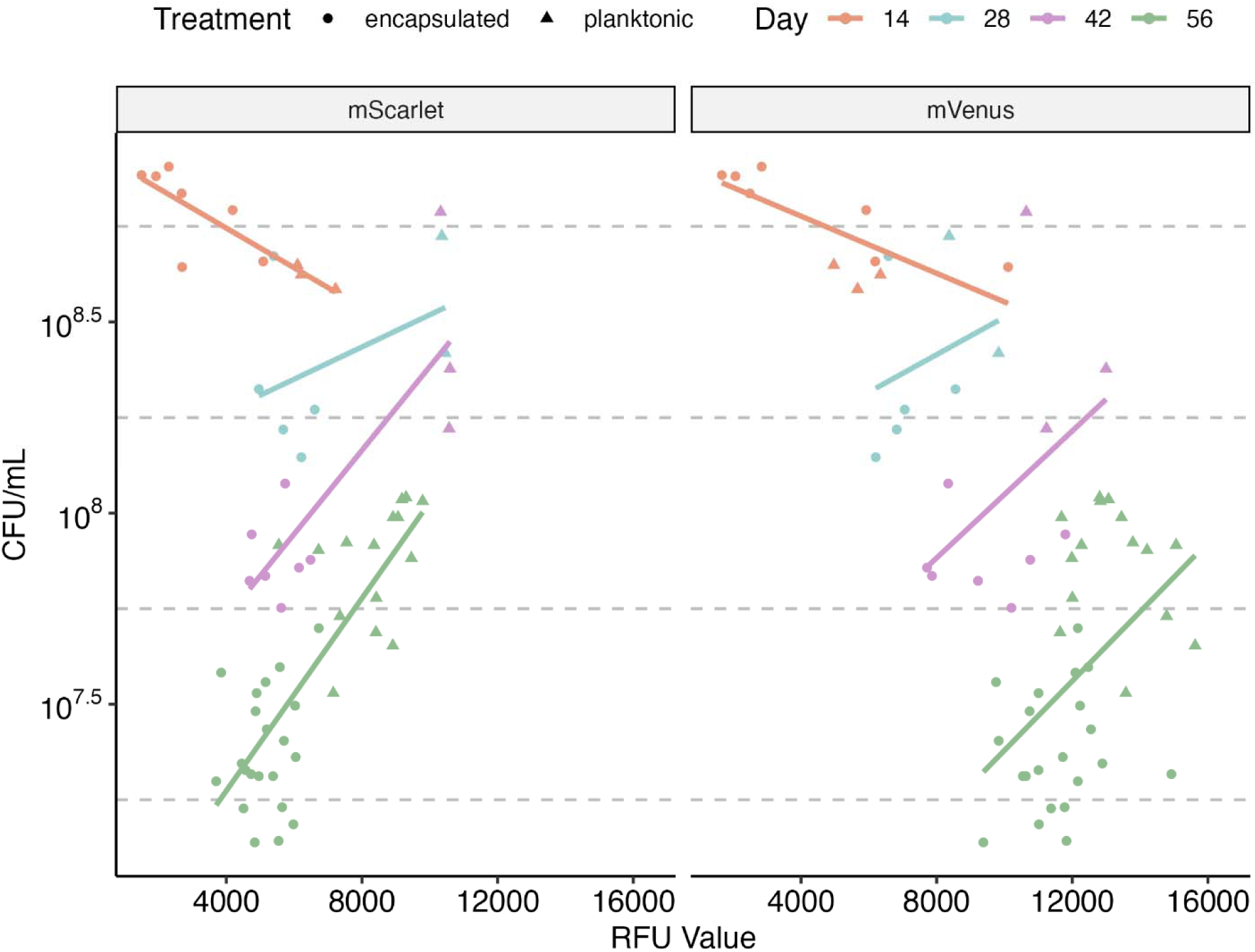
CFU/mL versus relative fluorescence units (RFU) for *P. putida* in supernatant throughout experiment. Linear line of best fit is shown for each day.

Multiple linear regression models were used to evaluate the contributions of time, treatment, and fluorescence to supernatant cell concentration (Table S1). Together, these analyses demonstrate that fluorescence alone was insufficient to predict viable cell concentration and that its relationship with CFU/mL depended strongly on time.

## Discussion

This study demonstrates that the effects of alginate encapsulation on bacterial growth and persistence are not generalizable but depend on bacterial species, nutrient environment, and capsule formulation. Encapsulation had relatively little effect on the long-term persistence of *P. putida* under the conditions tested, whereas *N. aromaticivorans* exhibited substantially greater long-term abundance when encapsulated than when grown planktonically. Short-term growth responses were similarly context dependent, with the effects of encapsulation differing between LB and sRB15 and between the two bacterial species. These findings are consistent with previous studies showing that immobilization can enhance bacterial growth and cell density (Garcia et al., 2019; Qi et al., 2006), but further demonstrate that these benefits depend on interactions among the microorganism, capsule properties, and environmental conditions. Potential mechanisms underlying improved performance of encapsulated cells include localized nutrient availability and protection from environmental stress (Sun et al., 2007). Together, these results highlight the importance of considering both microbial physiology and site-specific environmental conditions when designing encapsulated inoculants for bioaugmentation applications.

The two bacterial species exhibited notably different responses to encapsulation. *P. putida* grew robustly under both encapsulated and planktonic conditions, with the primary short-term benefit of encapsulation observed as an increased maximum cell density in sRB15 with chitosan-coated capsules. Although encapsulation decreased the initial growth rate and overall growth of *P. putida* in LB, all treatments ultimately achieved similar maximum densities, and long-term total reactor concentrations did not differ between encapsulated and planktonic conditions. Thus, under the relatively simple conditions evaluated here, encapsulation provided little long-term advantage for maintaining *P. putida*. However, these experiments did not include microbial competition or many of the environmental stresses likely to occur in contaminated environments. Such conditions could increase the relative advantage provided by encapsulation.

In contrast, *N. aromaticivorans* exhibited a stronger and more environment-dependent response to encapsulation. Encapsulation increased maximum cell density and overall growth in nutrient-rich LB but decreased initial growth rate and maximum density in sRB15. Reduced growth under lower-nutrient conditions could reflect mass-transfer limitations within the capsule, as diffusion through immobilization matrices can constrain nutrient availability (Jeong & Irudayaraj, 2023; Sønderholm et al., 2017). Despite this reduction in growth rate, encapsulation substantially shortened the lag phase of *N. aromaticivorans* in sRB15. One potential explanation is that immobilization creates a higher effective local inoculum, which can influence lag time (Ardré et al., 2022; Augustin et al., 2000). Shorter lag phases have similarly been reported for microorganisms immobilized in alginate beads and on carbon fiber (Boshagh et al., 2019; Ercole et al., 2021). Under nutrient-rich conditions, the increased maximum density of encapsulated *N. aromaticivorans* may also reflect improved resource use or enhanced local signaling among immobilized cells (Li et al., 2023).

These short-term responses were complemented by a pronounced long-term benefit of encapsulation for *N. aromaticivorans*. After 8 weeks in sRB15, total *N. aromaticivorans* concentrations were substantially higher in encapsulated reactors than under planktonic conditions, whereas no comparable encapsulation benefit was observed for *P. putida*. The relatively limited release of *N. aromaticivorans* into the surrounding medium may reflect slower growth within the capsules under low-nutrient conditions. Slower cell proliferation could reduce physical pressure on the alginate matrix and consequently decrease capsule disruption and cell release (Čejková et al., 2013; Stewart & Robertson, 1989). More broadly, these findings are consistent with previous studies demonstrating that the effects of encapsulation on viability are strain dependent (Borges et al., 2012; Sousa et al., 2012; Yeung et al., 2016).

Differences in ecological growth strategy may help explain the contrasting responses of the two organisms. *N. aromaticivorans* has fewer 16S rRNA gene copies than *P. putida* (3 versus 6, respectively), a trait generally associated with slower growth and greater resource-use efficiency (Klappenbach et al., 2000; Lauro et al., 2009; Roller et al., 2016). Thus, *N. aromaticivorans* exhibits traits more characteristic of an oligotrophic growth strategy, whereas *P. putida* exhibits traits associated with faster-growing copiotrophs (Lauro et al., 2009). Encapsulation may therefore provide a greater persistence advantage to relatively slow-growing organisms that would otherwise be less competitive under planktonic conditions. This could be particularly valuable for bioaugmentation because organisms with resource-efficient growth strategies may be well suited for sustained degradation of recalcitrant contaminants under resource-limited or stressful conditions (Brzeszcz et al., 2016; Thompson et al., 2005). Communities enriched in these ecological strategies have also been associated with increased PAH degradation and degradative plasmid conjugation (Varner et al., 2022).

Capsule formulation also influenced bacterial behavior, although changes that benefited short-term growth did not necessarily improve long-term persistence. Chitosan coating increased some short-term growth metrics, including maximum cell density, but resulted in lower cell concentrations within capsules after 8 weeks. Chitosan can exhibit antimicrobial activity depending on concentration and environmental conditions (Erdélyi et al., 2022; Klinkenberg et al., 2001; N. Liu et al., 2006; Tayel et al., 2010), with its antimicrobial properties also influenced by pH and solubility (Barreto et al., 2010; Rabea et al., 2003). Alternatively, the coating process itself can alter alginate matrix properties. Chitosan treatment has been reported to cause swelling and mechanical weakening of alginate gels, potentially facilitating cell release (Ta et al., 2021), while chitosan membranes can also restrict capsule pores and nutrient transport (Ji et al., 2019). Wen-tao et al. (2005) similarly reported that chitosan membrane formation decreased cell leakage and suggested that the membrane acted as a barrier to nutrient transport. These competing effects may explain why chitosan produced some apparent short-term growth advantages but no clear long-term benefit in the present study.

Alginate concentration also affected the distribution of viable cells within the encapsulated system, although it did not significantly alter total reactor concentrations after 8 weeks. This distinction between capsule-associated and total cell abundance is important for designing bioaugmentation systems because the desired formulation may depend on whether the application requires prolonged retention of biomass within a delivery matrix or more rapid release of cells into the surrounding environment. Similarly, initial cell loading did not significantly affect long-term cell concentrations under the conditions evaluated here, suggesting that increasing the initial inoculum within the range tested may not necessarily improve long-term persistence. Collectively, these results indicate that capsule formulation should be selected based on the desired balance among growth, retention, and release rather than assuming that increased polymer concentration, coating, or initial cell density will universally improve performance.

The biphasic growth profiles observed for some encapsulated treatments further illustrate the difficulty of distinguishing growth within capsules from growth of cells released into the surrounding medium. Although alginate utilization could theoretically contribute to multiple growth phases, neither organism grew when alginate was supplied as the sole carbon source (Fig. S3), making diauxic growth on alginate unlikely (Rojas-Padilla et al., 2022). Instead, the observed profiles may reflect an initial release of cells from the capsules followed by progressive growth of released cells in the surrounding medium (Corbo et al., 2011; Kaur et al., 2023). Once released cells begin growing exponentially, their contribution to bulk optical density can dominate over continued release from the capsule (Suzuki et al., 1998). Consequently, optical-density measurements of encapsulated cultures should be interpreted cautiously because they do not independently quantify growth within the capsule and surrounding medium.

Accurate enumeration of encapsulated cells presents a related methodological challenge. Sodium citrate is widely used to reverse alginate crosslinking and release immobilized biomass for enumeration (Feijoo-Siota et al., 2008; Partovinia & Rasekh, 2018; Sachan et al., 2009; Sohail et al., 2013; Sousa et al., 2012; Yao et al., 2018) and has been described as a standard approach for enumerating immobilized biomass (Serp et al., 2000). However, sodium citrate can also exhibit antimicrobial effects (Khayat et al., 2022; Nagaoka et al., 2010). Consistent with these reports, prolonged sodium citrate exposure decreased recoverable *P. putida* in our experiments, emphasizing that decapsulation conditions can influence apparent viability. The 15-min exposure used here provided sufficient capsule dissolution while minimizing loss of culturability. This methodological consideration is important when comparing the viability of encapsulated microorganisms across studies using different decapsulation procedures.

Fluorescence provided an additional means of tracking encapsulated cells but was not a reliable time-independent proxy for viable cell abundance. Fluorescence has previously been correlated with bacterial cell counts and used as a convenient alternative to optical density for monitoring growth (Schlechter et al., 2021; Wilson et al., 2018). Fluorescently tagged or stained microorganisms have also been used to examine growth, survival, and cell abundance within encapsulation matrices (Moore et al., 2015; Nandy et al., 2021; Podrazky & Kuncova, 2005; Sønderholm et al., 2017). Although fluorescence and viable counts were strongly related in the initial standard curves, fluorescence alone was a poor predictor of viable *P. putida* concentrations over the 8-week experiment. Incorporating time substantially improved this relationship, indicating that fluorescence per culturable cell changed over the course of the experiment. Potential explanations include changes in cellular physiology, fluorescent protein expression and maturation, and growth rate (Leveau & Lindow, 2001), as well as persistence of fluorescent proteins after reductions in culturable cell abundance (Snapp, 2009). In addition, stressed or viable-but-nonculturable cells may retain fluorescence despite not being recovered by plate counts (Lowder et al., 2000; Steff et al., 2001). Thus, fluorescence can complement culture-based measurements but should be independently validated when used to estimate viable biomass over extended periods.

Overall, these findings demonstrate that there is unlikely to be a single optimal alginate encapsulation formulation across microbial inoculants and environmental conditions. Rather, the benefits of encapsulation depend on interactions among microbial physiology, nutrient availability, and capsule properties. Encapsulation provided a particularly strong long-term persistence advantage for *N. aromaticivorans*, whereas *P. putida* maintained high abundance even without encapsulation under the relatively simple conditions tested. Capsule composition further influenced the balance between cell retention and release, emphasizing that formulation should be matched to the intended function of the inoculant. Although this study focused on bacterial growth and persistence rather than directly measuring contaminant degradation, these findings provide design principles for selecting and formulating encapsulated inoculants for bioaugmentation. Future work should evaluate these relationships in more environmentally complex systems that include indigenous microbial competition, heterogeneous nutrient availability, and contaminant exposure, while directly linking inoculant persistence and release to contaminant degradation.

## Acknowledgements

The authors thank the Duke Superfund Data Management & Analysis Core for their guidance on data analysis.

## Funding Sources

Research reported in this publication was supported by the National Institute of Environmental Health Sciences of the National Institutes of Health under Award Number P42ES010356 (Duke University Superfund Research Center – Developmental Co-Exposures: Mechanisms, Outcomes, and Remediation). The content is solely the responsibility of the authors and does not necessarily represent the official views of the National Institutes of Health. This work was also supported by the National Science Foundation (NSF) GRFP under grant DGE 2139754.

## Author Contributions

AMF: Conceptualization, Data curation, Investigation, Methodology, Visualization, Writing – original draft, editing. CKG: Conceptualization, Funding acquisition, Resources, Supervision, Writing – review and editing

## Data Availability Statement

The data that support the findings of this study are publicly available in the Duke University Research Data Repository (doi: https://doi.org/10.7924/r4r564).

## Conflicts of Interest

The authors declare no conflicts of interest.

## Notes

https://doi.org/10.7924/r4r564

### Competing Interest Statement

The authors have declared no competing interest.

