## Supplemental Materials for "Effect of alginate encapsulation on growth and viability of polycyclic aromatic hydrocarbon-degrading bacteria varies by environment, species, and capsule design"

for

Supplementary Methods

**R analysis with gcplyr.** After raw OD data were blank-corrected, a small pseudocount (0.001) was added to avoid zeros. For smoothing, moving average filters were used (window width = 7 time points) for downstream analysis. Per-capita growth rates were calculated with calc_deriv with a window width of 3 time points on log-transformed smoothed OD data. For wells which contained too few data points before phase-1 maximum and gcplyr’s lag_time() failed, a manual tangent lag time was computed from the phase peak. The phase-shift time was defined as the first local minimum of the per-capita derivative occurring after the phase-1 maximum. Phase 2 maxima were identified as the first local maximum of the per-capita derivative after the phase-shift, with an expanded extrema search window (window width = 7) to prevent spurious maxima. Groups not expected to show two phases based on visual inspection of individual well data were explicitly excluded from phase-2 metric calculation. For area under the curve (AUC), strain-specific integration times were employed (113 h for *N. aromaticivorans* and 71 h for *P. putida*) since each strain required different lengths of time to reach stationary phase. Diagnostic plots (per-well smoothed OD, log OD, and per-capita derivatives) were generated to confirm the correct calculation of growth metrics (data not shown).

**Growth on sole carbon sources.** The ability of *P. putida* and *N. aromaticivorans* to grow on alginate or chitosan as a sole carbon source was evaluated through OD_600_ growth curves. Strains were prepared and adjusted in concentration as previously described, then washed 3x with PBS. 5 μL of cells were inoculated into 1000 μL of media in a 48-well plate containing the following media: sRB15 + no carbon (negative control), sRB15 + 0.2% alginate, sRB15 + 0.2% chitosan, sRB15+ pyruvate (positive control), and LB (positive control). The plate was incubated shaking (orbital, 3mm amplitude) at 30°C for 7 days. Blank media of each type was included to confirm the absence of contamination. Measured OD_600_ values were corrected based on initial OD_600_ values at time zero.

Supplemental Results

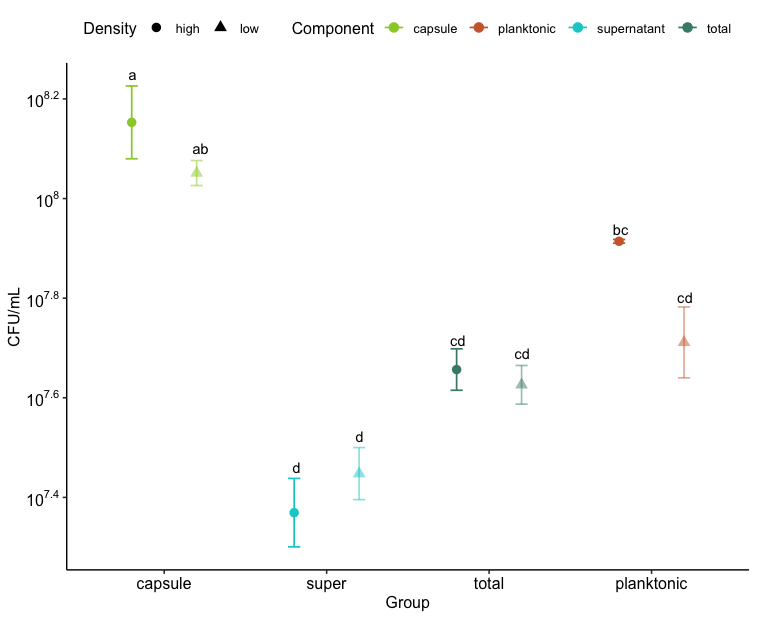

Figure S1. Viability/survival after 8-week incubation for *P. putida* treated with 10⁶ (low) or 10⁷ (high) CFU/mL initial cell densities in encapsulated and planktonic treatment groups. Means not sharing a letter are significantly different by Tukey's HSD test (*p* < 0.05). Error bars represent standard error (n = 3).

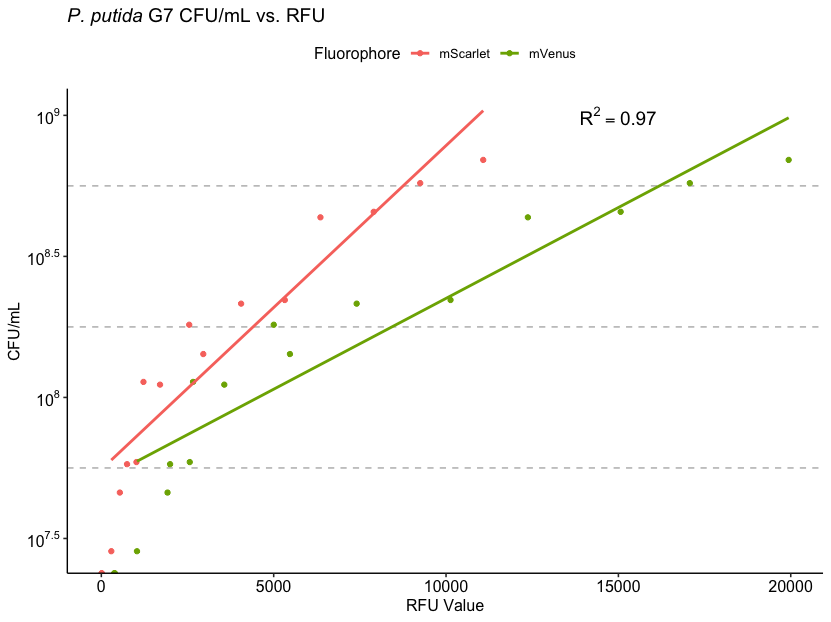

Figure S2. Cell concentration (CFU/mL) versus relative fluorescence unit (RFU) value for *P. putida* containing 2 fluorophores: mScarlet and mVenus. Linear line of best fit shown.

Table S1. Summary of regression analysis.

|  | Model 1 | Model 2 | Model 3 | Model 4 | Model 5 |
| --- | --- | --- | --- | --- | --- |
| Intercept | 7.93 *** | 7.80 *** | 7.93 *** | 7.77 *** | 7.83 *** |
|  | (0.03) | (0.03) | (0.04) | (0.04) | (0.04) |
| Day | -0.45 *** | -0.47 *** |  | -0.42 *** | -0.43 *** |
|  | (0.03) | (0.03) |  | (0.06) | (0.03) |
| Encapsulation_Treat |  | 0.37 *** |  | 0.38 *** | 0.17 * |
|  |  | (0.05) |  | (0.06) | (0.08) |
| mVenus |  |  | -0.46 *** | -0.07 |  |
|  |  |  | (0.05) | (0.07) |  |
| mScarlet |  |  | 0.23 *** |  | -0.12 * |
|  |  |  | (0.05) |  | (0.06) |
| mVenus:Day28 |  |  |  | -0.07 |  |
|  |  |  |  | (0.10) |  |
| mVenus:Day42 |  |  |  | 0.11 |  |
|  |  |  |  | (0.14) |  |
| mVenus:Day56 |  |  |  | 0.06 |  |
|  |  |  |  | (0.09) |  |
| mScarlet:Day28 |  |  |  |  | 0.17 |
|  |  |  |  |  | (0.09) |
| mScarlet:Day42 |  |  |  |  | 0.31 *** |
|  |  |  |  |  | (0.07) |
| mScarlet:Day56 |  |  |  |  | 0.31 *** |
|  |  |  |  |  | (0.07) |
| N | 63 | 63 | 63 | 63 | 63 |
| R^2^ | 0.74 | 0.86 | 0.59 | 0.87 | 0.91 |
| p | *** | *** | *** | *** | *** |
| AIC | 16.5 | -20.5 | 47.6 | -22.1 | -47.0 |
| All continuous predictors are mean-centered and scaled by 1 standard deviation. The outcome variable is in its original input units (log-transformed CFU/mL). Standard error is reported in parentheses below each coefficient estimate. *** p < 0.001; ** p < 0.01; * p < 0.05. | | | | | |

**Regression analysis.** Several iterations of the multiple regressions were carried out to best explain the data. The main dynamic we observed in the supernatant was a linear decay over time in log scale bacterial concentrations, shown in Model 1 (Table S1). The variable “Day” alone explains most of the variance in the data (R2 = 0.74). Encapsulation treatment (“Encapsulation_Treat”) is also statistically significant, suggesting that the two treatments led to different average concentrations. When accounting for day and treatment (Model 2), these variables explain 86% of the variance (R2 = 0.86).

Fluorescence values alone (Model 3) do not explain the data very well (R2 = 0.59); thus, fluorescence is not a strong predictor without accounting for other variables. In fact, in Model 3, the coefficient for mVenus is negative, though initial standard curves had suggested a positive relationship between fluorescence and concentration. However, when accounting for the interaction between fluorescence and time for mScarlet (Model 5), the model has a fit of R2 = 0.91, with the expected positive coefficients for fluorescence within each day. While the addition of mScarlet values improves the model, mVenus (Model 4) does not add explanatory value, likely because the mVenus measurements contained more noise and variation.

Akaike Information Criterion (AIC) indicates that Model 5, incorporating day, treatment, fluorescence, and interactions provides the best trade-off between fit and model complexity. Model 5’s AIC is substantially lower than Model 2, demonstrating that despite having more predictors, it provides a significantly better fit to the data, justifying the added complexity. The AIC shows that Model 3 is a poor model, reinforcing that fluorescence alone is an insufficient predictor.

Our fluorescence measurements on Day 14 present an inverse relationship between fluorescence and CFU/mL, in contrast with the rest of the measurements and the positive relationship demonstrated in the standard curves. One potential explanation for this inverse relationship is the inner filter effect; since the cell densities were high at this point in the incubation, excess cell density led to light absorption by the sample and reduced fluorescence intensity, which could be resolved by dilution (Surribas et al., 2006).

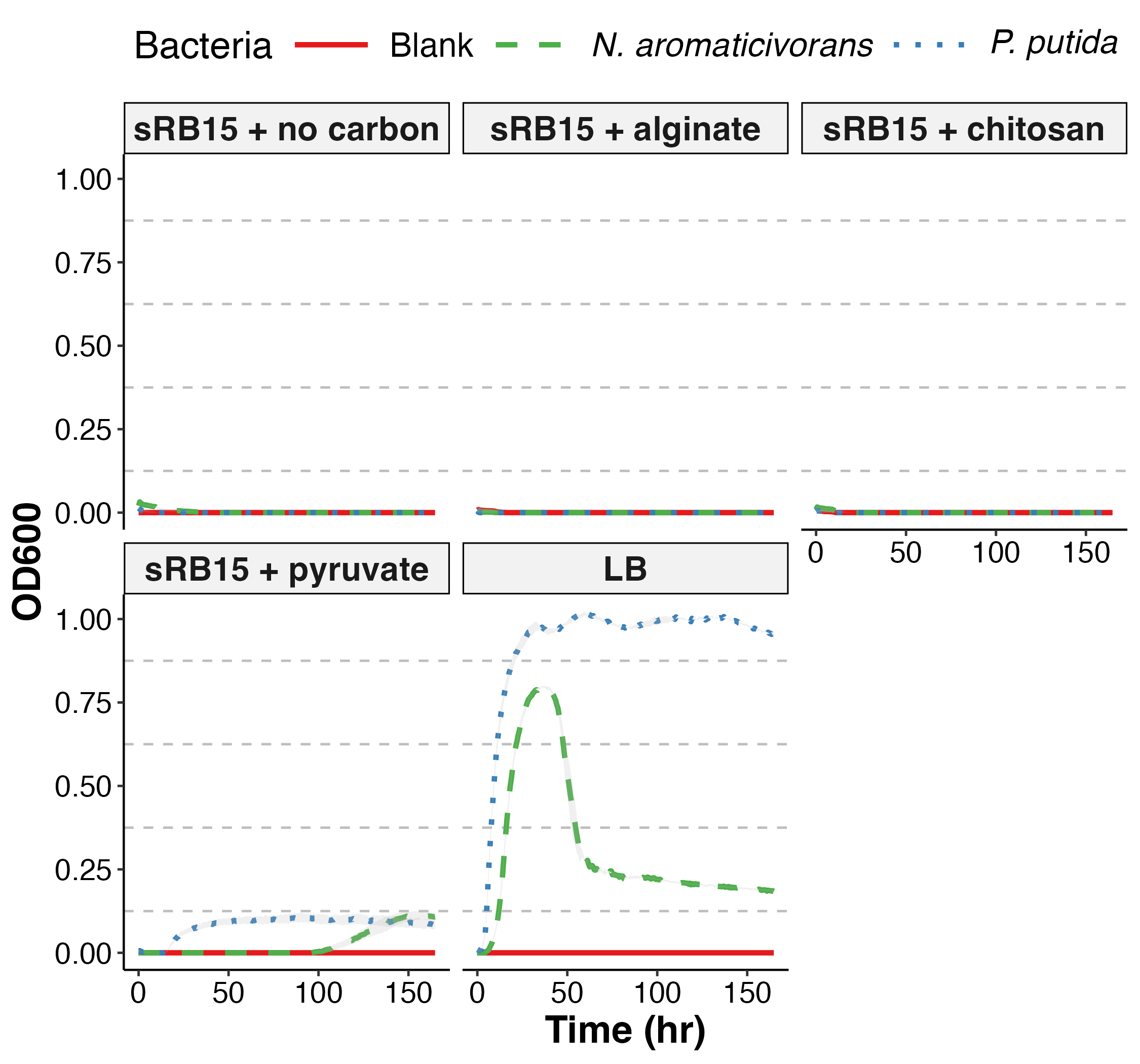

Figure S3. Growth curves for *P. putida* *and N. aromaticivorans* with different sole carbon sources (alginate, chitosan, and pyruvate) as well as a no carbon negative control and an LB positive control. Grey regions represent standard error (n = 3).
